# Nanoaperture fabrication via colloidal lithography for single molecule fluorescence imaging

**DOI:** 10.1101/596460

**Authors:** Ryan M. Jamiolkowski, Kevin Y Chen, Shane A. Fiorenza, Alyssa M. Tate, Shawn H. Pfeil, Yale E. Goldman

**Affiliations:** Pennsylvania Muscle Institute, Perelman School of Medicine, University of Pennsylvania, Philadelphia, Pennsylvania, United States; Department of Physics, West Chester University, West Chester, Pennsylvania, United States

## Abstract

In single molecule fluorescence studies, background emission from labeled substrates often restricts their concentrations to non-physiological nanomolar values. One approach to address this challenge is the use of zero-mode waveguides (ZMWs), nanoscale holes in a thin metal film that physically and optically confine the observation volume allowing much higher concentrations of fluorescent substrates. Standard fabrication of ZMWs utilizes slow and costly E-beam nano-lithography. Herein, ZMWs are made using a self-assembled mask of polystyrene microspheres, enabling fabrication of thousands of ZMWs in parallel without sophisticated equipment. Polystyrene 1 μm dia. microbeads self-assemble on a glass slide into a hexagonal array, forming a mask for the deposition of metallic posts in the inter-bead interstices. The width of those interstices (and subsequent posts) is adjusted within 100-300 nm by partially fusing the beads at the polystyrene glass transition temperature. The beads are dissolved in toluene, aluminum or gold cladding is deposited around the posts, and those are dissolved, leaving behind an array ZMWs. Parameter optimization and the performance of the ZMWs are presented. By using colloidal self-assembly, typical laboratories can make use of sub-wavelength ZMW technology avoiding the availability and expense of sophisticated clean-room environments and equipment.

## Introduction

Single molecule fluorescence techniques are valuable tools in biophysical research. By avoiding the averaging inherent in bulk measurements, they can distinguish subpopulations of molecules, directly observe the trajectory and timing of enzymatic reaction steps without needing to synchronize a population, and can enable study of rare events and conformational fluctuations [1]. These techniques include super-resolution microscopy to track single molecule motion, fluorescence resonance energy transfer (FRET) to detect nanometer-scale distance changes, and polarized total internal reflectance (polTIRF) microscopy that measures angular changes of a macromolecule [2–4]. All single-molecule fluorescence techniques require careful optimization of signal-to-noise ratio due to their inherently limited signal.

The two most widely adopted schemes for reducing fluorescence background from out of focus fluorescent probes are total internal reflection fluorescence (TIRF) microscopy and confocal microscopy. Both techniques create small optically defined volumes of detection. In the case of TIRF, incident light reaches the glass-water interface at an angle greater than the critical angle for total internal reflection and is reflected. This creates a near-field standing wave (the evanescent wave) on the aqueous side that decays exponentially with distance from the boundary. This localized oscillating electromagnetic field excites only the fluorescent molecules very close (<100 nm) to the glass-water interface, significantly reducing background fluorescence. In confocal microscopy, a confocal pinhole in a conjugate image plane is used in conjunction with a focused laser spot to define a femtoliter scale detection volume.

Despite the advantages offered by these arrangements, background fluorescence from labeled substrates in solution remains a technical challenge, requiring fluorescent substrates to be used at nano-/pico-molar concentrations. However, biological processes often occur at concentrations up to a million-fold greater (micro-/milli-molar). A potential solution is nanofabricated zero-mode waveguides (ZMWs) [5–7], arrays of holes (30-300 nm diameter) in a thin, opaque metal layer (typically aluminum, chromium, or gold) on a glass substrate [7]. These sub-wavelength apertures do not propagate optical modes in the visual spectrum, but illumination will create an evanescent wave, like in TIRF, that decays inside the well [8,9].

For fluorescence microscopy, ZMWs are most applicable when their size is sufficiently small that the excitation light decays within the waveguide (*λ* > *λ*_*c*_). By constraining the observation volume physically (in the plane of the slide) and optically (along the optical axis), ZMWs give observation volumes of atto-liter scale or smaller, allowing use of micro-/milli-molar fluorescent substrates without prohibitive background fluorescence [7]. At µM concentrations of fluorescent substrates, only one or a few probes are simultaneously within the ZMW, exchanging rapidly with the non-excited bulk phase.

ZMWs have been produced by directly patterning the metal layer using ion beam milling [10,11], via electron beam lithography and subsequent dry-etching [5,12], or metal lift-off [13–16]. Although the presence of thousands of wells on each substrate allow parallel collection of data during an experiment, each well of a ZMW array is either fabricated in series, a slow and expensive process, or in parallel with advanced deep-UV lithography [17]. In contrast to these top-down methods, which require significant overhead in the form of specialized equipment, nanosphere (or “colloidal”) lithography (NSL) uses the self-assembly of an ordered crystal to create a mask for nanopatterning.

NSL was first introduced nearly 40 years ago [18–20], and has been used to fabricate many different nanostructures, including dry-etched SiO_2_ nanoplates for polymer imprinting [21], large scale arrays of silicon nanowires [22], ordered arrays of gold nanoparticles for catalysis of ZnO nanowire formation [23], surface for the selctive imobilization of biomolecules [24,25], and sensors based on surface enhanced Raman spectroscopy (SERS)[26]. Complex geometries can be obtained with multiple overlapping layers of micron and nanoscale spheres [27], or by tilting the masks relative to the metalizing vapor beam to change the projection geometry [28]. Nonetheless, NSL remains less widely used than electron-beam or photolithography.

To take advantage of NSL’s simplicity, flexibility and low cost, a method of fabricating ZMWs through colloidal templating of polystyrene beads was developed, using the self-assembly of an ordered structure to pattern arrays of nanoscale wells. This technique can be used to fabricate ZMWs from many different metals, including the commonly-used aluminum [5,6,29] and gold [30]. Importantly, this technique requires few specialized fabrication tools, and should be accessible to many labs that perform single-molecule experiments.

## Results and Discussion

### Formation and Annealing of Polystyrene Bead Mask

ZMWs were fabricated using colloidal lithography according to the schematic diagram in Fig 1. Suspensions of 1 µm polystyrene beads in 1:400 (v/v) TritonX100:ethanol were pipetted onto the centers of the coverslips, which had been kept at 85-90% relative humidity (see Materials and Methods, S1 Fig). When spherical particles are partially immersed in a liquid on a horizontal substrate, the liquid meniscus between them is deformed, which causes attraction between adjacent spheres via surface tension. In addition, as solvent evaporates from those menisci, convective flux further drives the particles into an ordered phase[31]. Thus, the bead suspensions spread into approximately 2 cm circular puddles in which 2D hexagonal lattices (Fig 1b, 2a-d) self-assemble, with thin voids between domains having different lattice orientations (Fig 2c,d).

**Fig 1.**
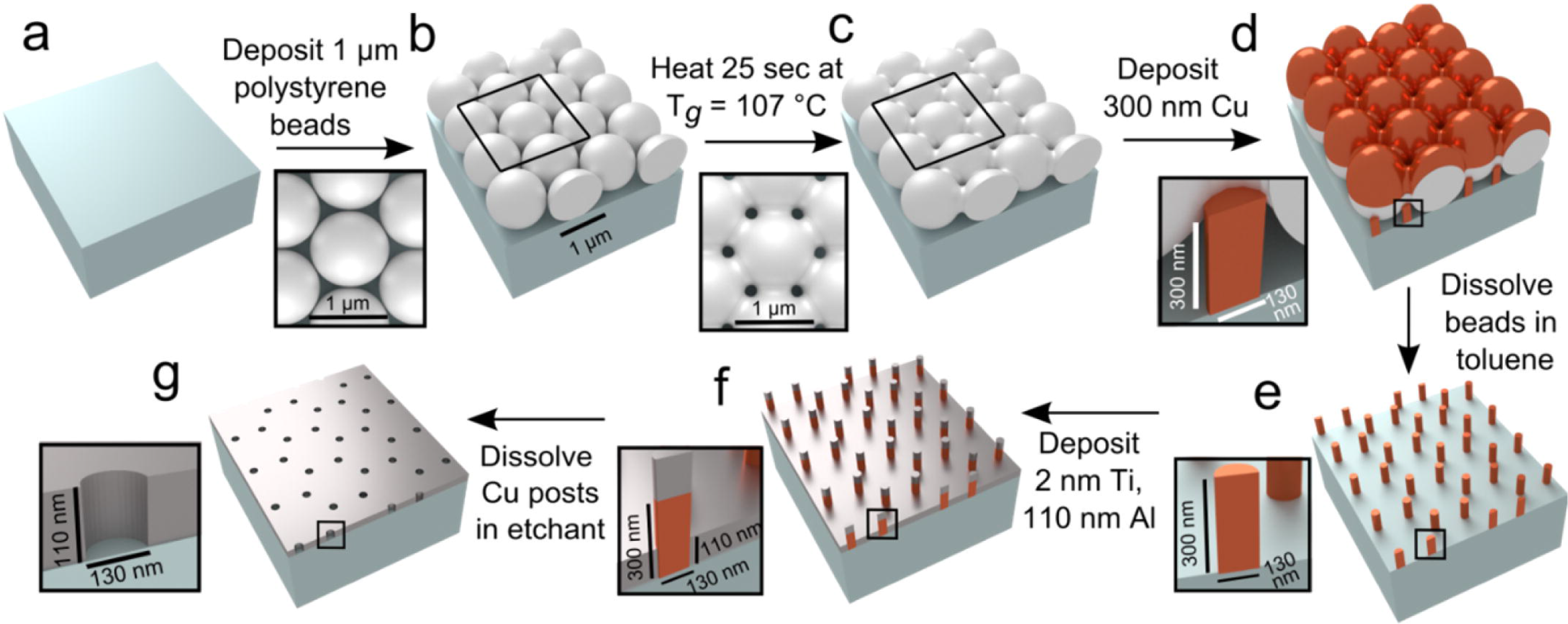
Schematic of aluminum ZMW fabrication. Black boxes represent expanded insets. T_*g*_ is the glass transition temperature.

**Fig 2.**
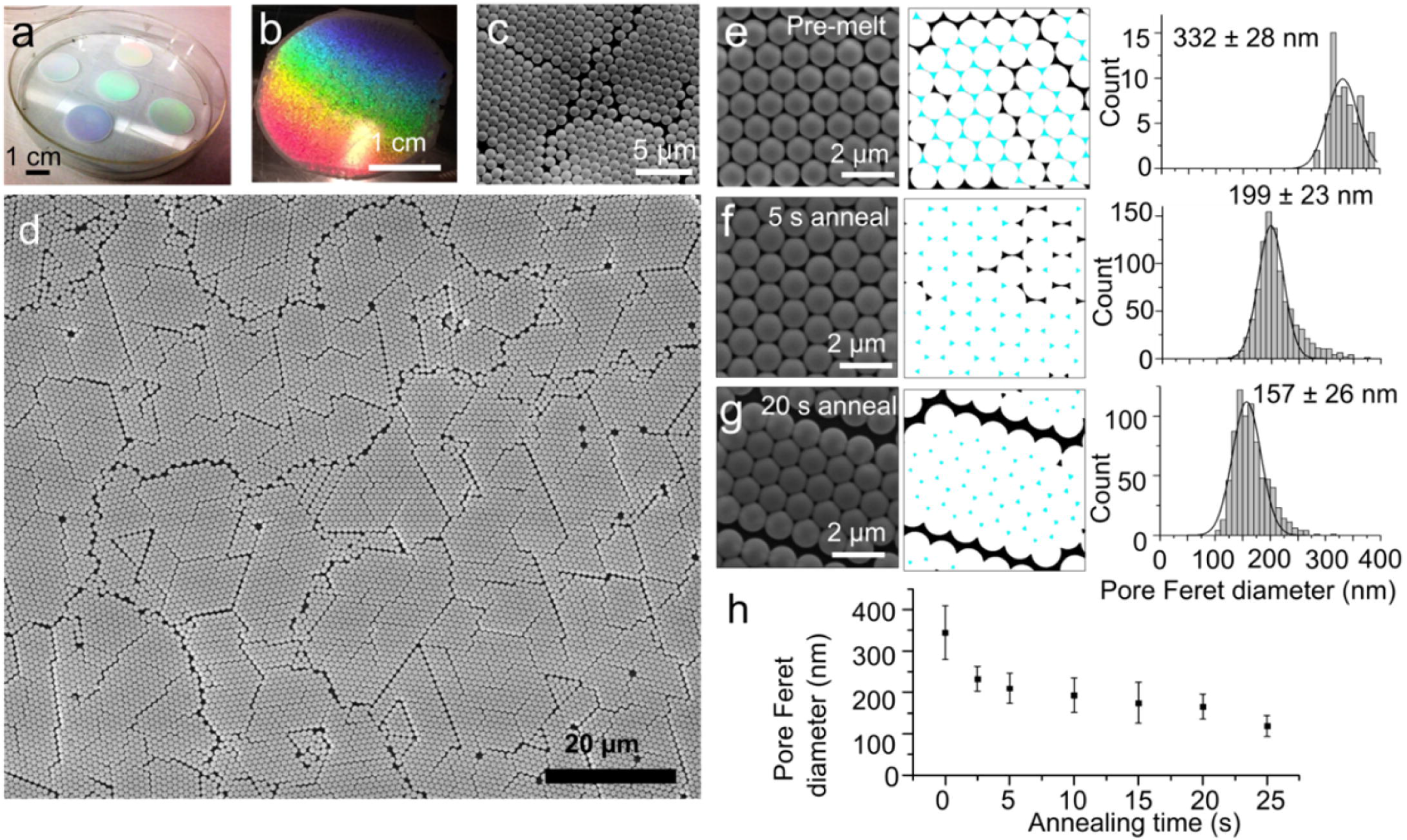
Annealing of 2D crystalline arrays of polystyrene beads. Macroscopic images of 2D crystalline arrays of polystyrene beads displaying rainbow structural coloration **(a,b)** and SEM images of those hexagonal arrays **(c,d).** SEM images of beads that were not heated **(e),** or were heated at 107 °C for 5 s **(f)** or 20 s **(g),** with corresponding interstitial areas selected (cyan) and Feret diameter histograms for the measured areas. **(h)** Interstitial Feret diameter from SEM images as a function of melting time. Error bars represent standard deviation.

The large size of these grains (~5-15 µm across, Fig 2c,d) uniformly covering 2 cm areas may represent a useful feature not only for the fabrication of ZMWs, but possibly for the fabrication of materials for batteries and energy storage [32], as well as nanowires for semiconductor devices [33]. The use of ethanol rather than water for the colloidal suspension differentiates this procedure from other published methods [34], allowing the evaporation to occur in only 2-3 minutes rather than two hours. Relative to other methods [34], this more robust deposition process only requires precise control of relative humidity, rather than of temperature, and surface tilt (as required by other methods).

When the lattice has properly assembled, the periodic surface of the bead array causes wavelength-dependent interference in reflected incident light and a brilliant, rainbow iridescence (Fig 2b) similar to that of peacock feathers and butterfly wings [35,36]. This striking structural coloration allows an immediate, macroscopic assessment of 2D lattice formation.

Since the interstices between the beads ultimately determine the cross-sectional shape and size of the ZMWs, the wells were made narrower and more cylindrical by briefly heating the polystyrene beads close to their glass transition temperature (approximately 107 °C) [37], allowing them to fuse with one another at their contact points (Fig 1c). Similar results have previously been obtained using microwave pulses [38,39], but the use of a standard laboratory hot plate simplifies the process. This procedure also partially decouples the size of the holes in the polystyrene mask from the size of the spheres used, which controls the ZMW spacing. Simply substituting smaller beads for the 1 μm ones would create smaller interstitial gaps, but would also reduce the spacing between the wells and prevent them from being completely resolved in optical images. The timing of this heating step needs careful adjustment. For a range of treatment times from zero to 30 seconds (at which duration the beads fuse together completely), there is a reproducible relationship between melting time and size. The resulting pores can be tailored to produce diameters in the range of 350-100 nm (Fig 2h), which is essential for constructing waveguides with cutoff wavelengths in the visible range appropriate for experiments with the most commonly used probes for single molecule fluorescence. In addition, a round cross-section is also important; in non-centro-symmetric waveguides, the transmission is polarization sensitive, which could compromise the attenuation of the ZMWs [41] or cause their fluorescence to be sensitive to the orientation of the macromolecules [42].

### Fabrication of ZMWs

#### Pore Creation

The subsequent text will primarily focus on the fabrication of aluminum ZMWs, which starts with the deposition of copper posts (Fig 1). Where relevant, we also note the differences in the procedure used to fabricate gold ZMWs, which starts with the deposition of aluminum posts (S2 Fig.).

After the hexagonal lattice of beads was formed and annealed, the resulting polystyrene masks were used during line-of-sight thermal or e-beam evaporative plating of 300 nm of copper or aluminum that reached the glass surface only through the interstices between the beads (Fig 1d, S2d). The mirrored top surfaces of the beads also enhanced the structural coloration, giving a vibrant rainbow appearance (Fig 3a, S3a).

**Fig 3.**
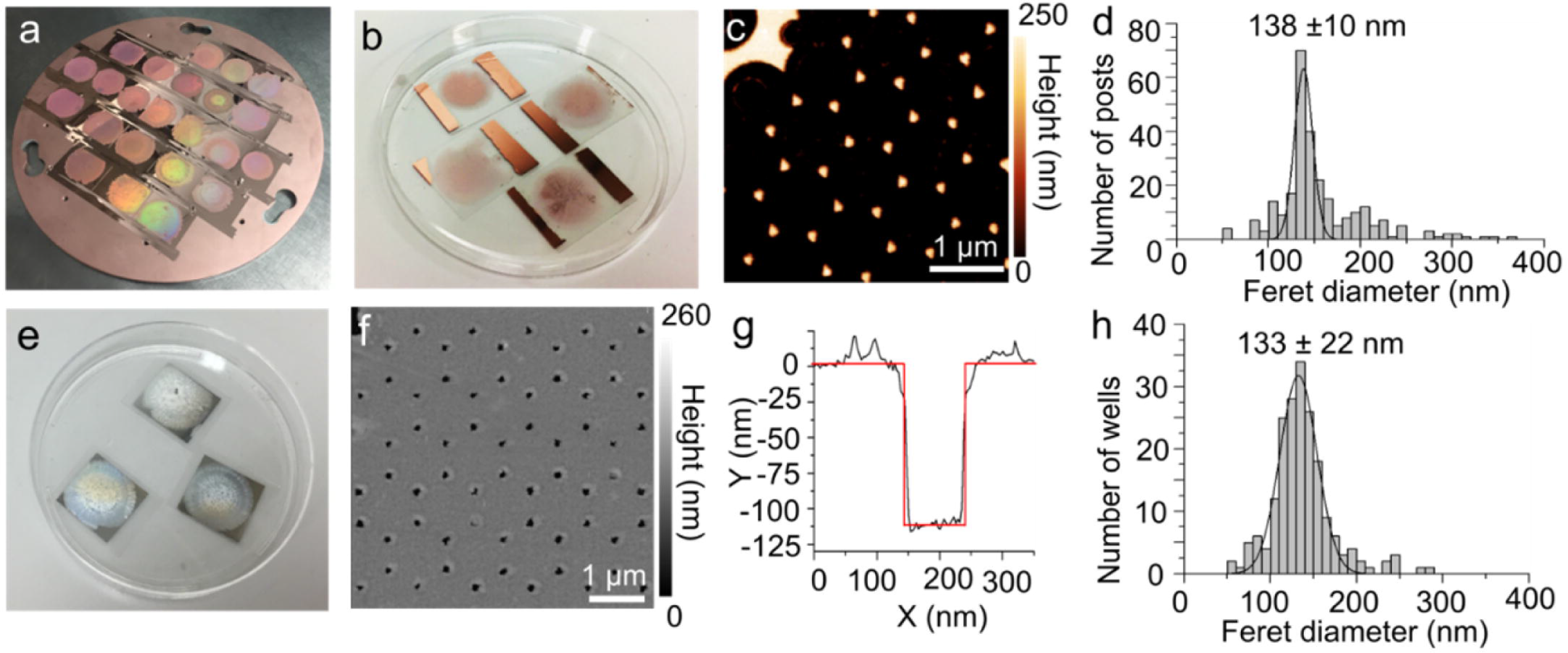
AFM Characterization of Al devices. **(a)** Structural coloration from the bead diffraction pattern following Cu evaporation. **(b)** Coverslips with centers patterned by array of copper posts following dissolution of polystyrene beads by toluene and tape pull of copper that had coated the tops of the beads. **(c)** AFM image of Cu posts on coverslip. **(d)** Distribution of Feret diameter of posts formed using a bead template annealed for 20 s. **(e)** Completed Al ZMWs. **(f)** AFM of ZMW wells formed after dissolving the narrow Cu posts. **(g)** AFM profile of depth of individual well. **(h)** Distribution of Feret diameter of wells in Al ZMW.

Removing the polystyrene mask, and the metal deposited onto it was accomplished by soaking in toluene, exfoliating the layer of metal on top of the softened beads with Scotch tape, and soaking again in toluene to dissolve any polystyrene remnants. This procedure left a hexagonal array of copper posts (Fig 1e, Fig 3b,c and Materials and Methods) or aluminum posts (Fig S2e, Fig S3b,c) with heights of 250-300 nm measured by atomic force microscopy (AFM),(Fig 3c, S3b). The maximum Feret diameters of the resulting posts have distributions centered at 130-140 nm (Fig 3d and S3d).

A cladding, 110 nm of aluminum or gold, was deposited around the posts (Fig 1f, S2f), which were selectively dissolved in the appropriate etchant (copper or aluminum) to leave arrays of pores to act as ZMWs (Fig 1g, 3e, S2g, S3e). The most efficient post dissolution occurred when the samples were removed from etchant after an hour, buffed with lens paper to disrupt any remaining posts, and placed back in the etchant for an additional hour (see Materials and Methods). AFM confirmed that most of the posts were dissolved and demonstrated that the wells were approximately 110 nm deep (Fig 3g), as expected. The distribution of Feret diameters of the wells was centered at ~130-140 nm (Fig 3h, S3h), with wider population distributions than at the post stage, since each successive processing step introduced some variability. In summary, the main population of wells were approximately 135 nm wide and 110 nm deep, similar to the dimensions of gold [30] and aluminum [29] ZMWs commonly made by other means.

### ZMW Functionalization

After these aluminum or gold ZMW arrays were fabricated, their surfaces were passivated and functionalized for use in single-molecule fluorescence experiments using protocols reported earlier for devices made by e-beam lithography. The aluminum ZMW surfaces were passivated with poly(vinylphosphonic acid) (PVPA) to block non-specific binding [5,6,29], while the gold ZMW surfaces were passivated with a self-assembled monolayer of methoxy-terminated, thiol-polyethylene glycol (PEG).[30] For both types, the glass surface at the bottom of the wells was functionalized with a chemically-orthogonal 16:1 mixture of methoxy- and biotin-terminated silane-PEG [43].

## Device Characterization

### In Situ Transmission Measurements

The faint transmission of light through the ZMWs when back-illuminated with sufficient intensity, using the illumination tower of the inverted microscope (Fig 4) allowed rapid initial characterization of the optical properties of the guides.

**Fig 4.**
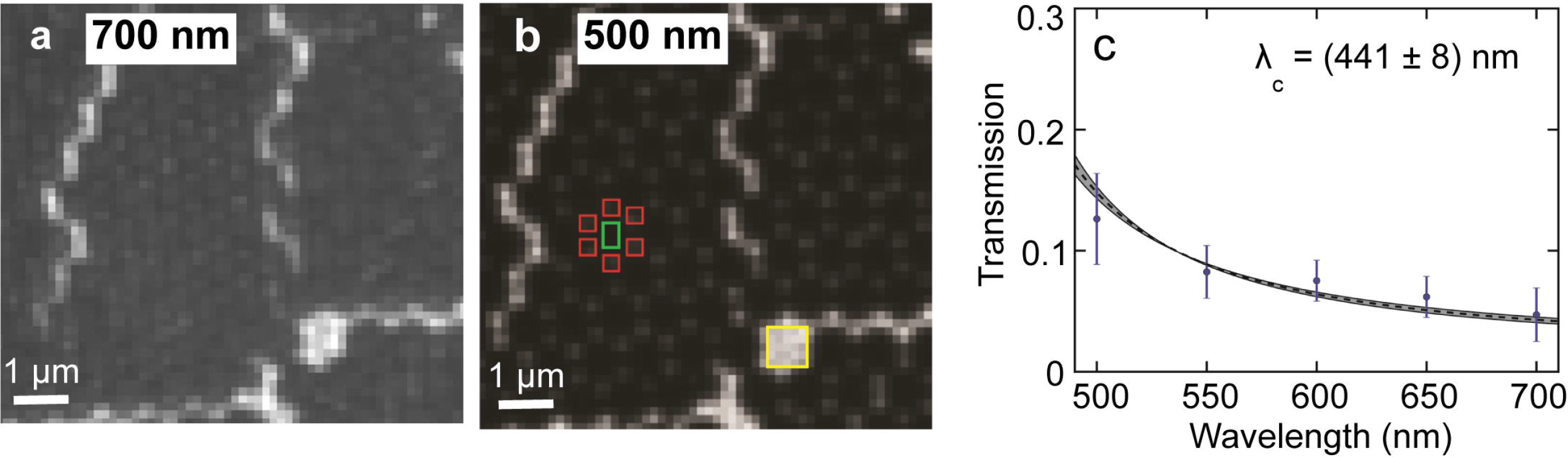
Transmission from back-illuminated guides. **(a)** Guides back illuminated with 700 nm light **(b)** Guides backlit with 500 nm light. Intensity data is collected from a set of pixels 2×2 in area centered on the guides (red boxes). The background level is collected from a box centered in the dark region between guides (green box). Unattenuated light intensity is collected from the closest large defect(yellow box). **(c)** Transmission data is fit to the model function 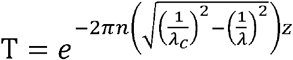, grey ribbon of values corresponds to the uncertainty in the guide thickness z.

To assess the optical attenuation of the ZMWs, we measured the transmission of the guides at 50 nm wavelength intervals from 500-700 nm (see Materials and Methods for details). As expected, transmission decreased as wavelength increased (Fig 4c). For the wavelength range measured, transmission through the wells was always <13% of the transmission through bare glass. This data was fitted using a simple model [44] for the transmission of a cylindrical waveguide (Fig 4c and Materials and Methods) which yielded a cut-off wavelength λ_c_ = 441 ± 8 nm. The consistency of the transmission data with this simple model provided an efficient initial quality control test.

Additionally, the unique pattern of bright defects caused by grain boundaries in the colloidal crystal mask was used to register the image coordinates of each ZMW (1500-2000 per field of view). These built in registration markers allowed stage drift to be compensated, and images to be registered across spectral channels.

## Single-Molecule Fluorescence Performance

### ZMW Performance Characterization with Labeled DNA

Wells were loaded with a biotinylated double stranded (ds) oligoDNA with Cy3 and Cy5 on opposite strands in positions known to give a FRET efficiency of ~0.7 using TIRF microscopy [45]. Before adding the oligo, the surface was first treated with 20 μM unlabeled streptavidin, which links the biotin on the PEG to that on the oligo using two of its four binding sites (see Materials and Methods). After incubating a 100 pM concentration of labeled molecules for 10 minutes, a substantial number of wells contained single fluorescent DNA duplexes, as judged by single step-wise photobleaching. For both aluminum ZMWs (Fig S4a) and gold ZMWs (S4c Fig), single molecule smFRET traces were detected from the biotin, Cy3, Cy5-dsDNA oligos in the wells under illumination with a 532 nm laser, that gave FRET efficiencies of ~0.7 as expected (S4b,d Fig).

The purpose of ZMWs is to attenuate background fluorescence intensity in the presence of a high concentration of labeled substrate diffusing freely in solution. In a typical TIRF setup without ZMWs, single-molecule traces cannot be detected above background fluorescence at probe concentrations above ~50 nM. After immobilizing biotin, Cy3-dsDNA oligos in ZMW wells and illuminating them with a 532 nm laser, however, we were able to detect individual immobilized fluorophores even at bulk concentrations of non-biotinylated Cy3-dsDNA as high as 1 μM (Fig 5a-d).

**Fig 5.**
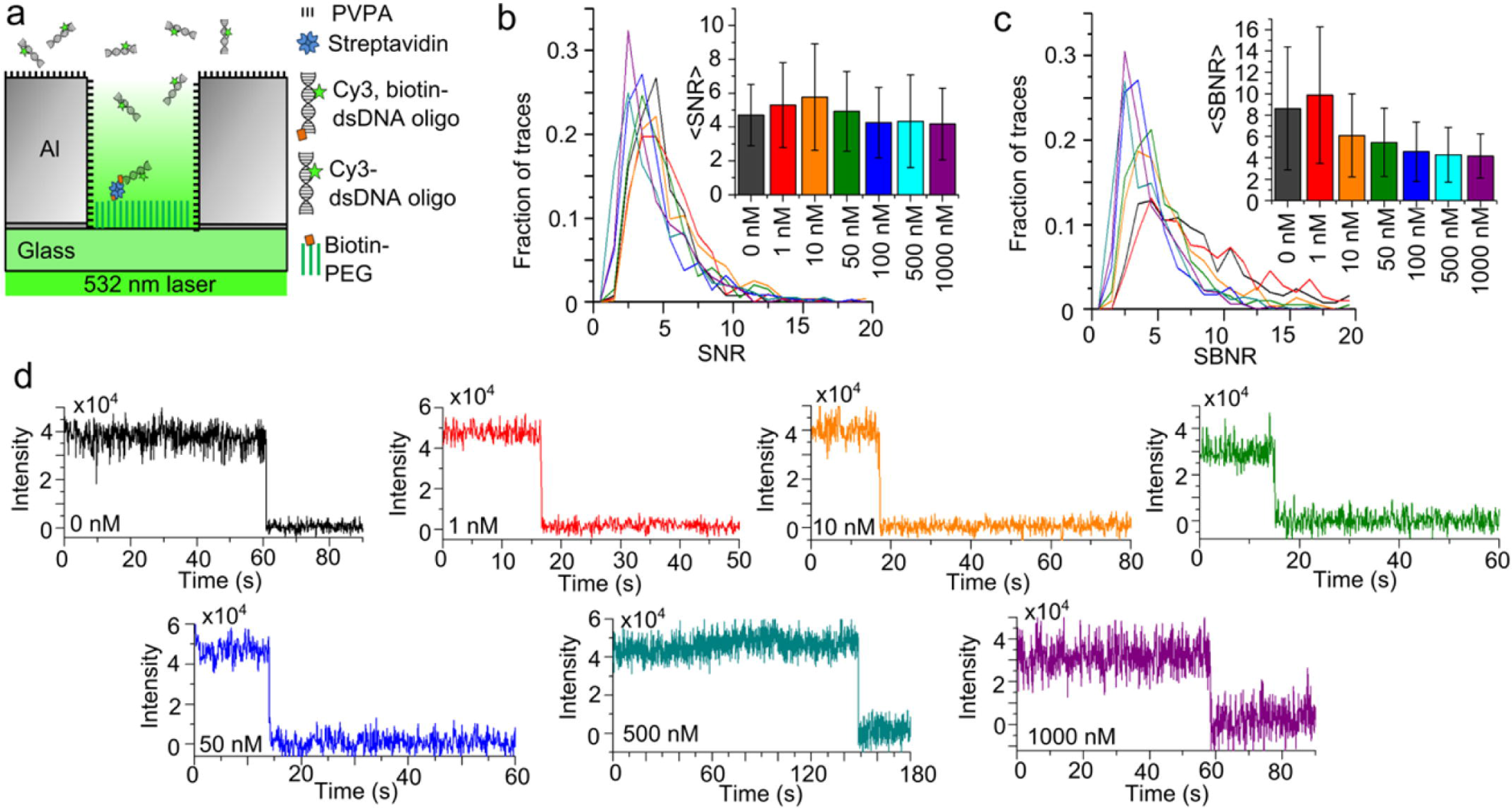
Single-Molecule performance with dsDNA at 532 nm excitation. **(a)** Schematic cartoon of biotinylated Cy3 dsDNA duplex immobilized in an Al ZMW while illuminated by 532 nm laser for direct excitation of Cy3, in the presence of non-biotinylated Cy3 dsDNA duplex in the solution. **(b)** Line histograms depict the distributions of the SNR and **(c)** SBNR of Cy3 emission from immobilized DNA oligos under direct excitation at different concentrations of background Cy3 dsDNA duplexes, with bar charts indicating the mean ± SD. **(d)** Example traces of Cy3 emission from immobilized oligos at bulk concentrations of Cy3 dsDNA in the surrounding solution of 0 nM (n = 262), 1 nM (437), 10 nM (515), 50 nM (195), 100 nM (346), 500 nM (349), 1000 nM (640).

The signal-to-noise ratios (SNRs) of those traces were calculated as the change in Cy3 fluorescence intensity due to photobleaching divided by the standard deviation of the Cy3 intensity before bleaching. The SNR did not vary significantly across all concentrations tested, with similar distributions (Fig 5b). The signal-to-background noise ratios (SBNRs) of those traces was calculated as the change in Cy3 fluorescence intensity due to photobleaching divided by the standard deviation of the intensity after the bleach, which is pure background fluorescence. The SBNR diminished slightly with increased concentration of labeled molecules in the solution (Fig 5c). However, it remained possible to record clear single molecule bleaches even at 1 μM (Fig 5d), which would not be possible in a confocal or TIRF microscope without the use of the ZMWs.

The ability to record single molecule fluorophores despite high concentrations of labeled substrate in the background was also observed for Cy5. Biotin, Cy3, Cy5-dsDNA oligos were immobilized in ZMWs and illuminated by a 640 nm laser (which directly excites only the Cy5) in the presence of background concentrations of non-biotinylated Cy5-dsDNA oligos as high as 1 μM (S5 Fig).

## tRNA-tRNA FRET in ZMW-Immobilized Ribosomes

Ribosomes programmed with a biotinylated mRNA were immobilized in the wells of Al ZMW arrays. PRE-translocation complexes were formed with Phe-tRNA^Phe^(Cy5) in the ribosomal P-site, and Val-tRNA^Val^(Cy3) in the A-site, where they exhibited FRET from Cy3 to Cy5 (Fig 6a). The pre-translocation complexes were stalled in the absence of the translocase EF-G·GTP, a condition that is known to allow transitions between “classical” and “hybrid” tRNA positions having high and moderate FRET efficiency[46,47]. In the presence of 500 nM free Phe-tRNA^Phe^(Cy5)·EF-Tu·GTP ternary complexes, smFRET traces showed ribosomes transitioning between a low FRET efficiency (~0.4) hybrid conformation and a high FRET efficiency (~0.7) classical conformation (Fig 6b) as previously observed[46,47]. Thus, these ZMWs fabricated via colloidal lithography allow single molecule fluorescence study of macromolecular dynamics at free concentrations of labeled substrates that would not be possible to approach using TIRF microscopy.

**Fig 6:**
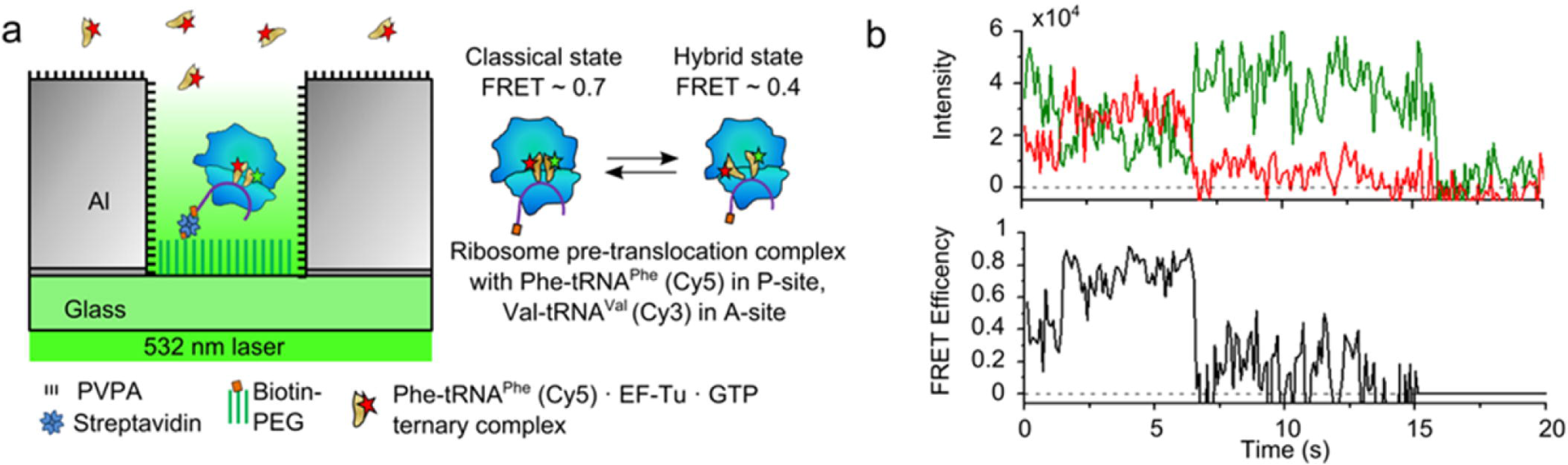
Application of devices to tRNA-tRNA FRET. **(a)** Schematic cartoon of ribosomes immobilized in an Al ZMW well via biotinylated mRNA, with tRNA-tRNA FRET between Phe-tRNA^Phe^(Cy5) in the ribosomal P-site and Val-tRNA^Val^(Cy3) in the A-site. The ribosomes were illuminated by 532 nm laser light, in the presence of 500 nM Phe-tRNA^Phe^(Cy5)·GTP·elongation factor-Tu (ternary complex) in solution. **(b)** Example FRET trace from a ribosome transitioning from low FRET hybrid state to high FRET classical PRE-translocation state before photobleaching of Cy5 at 7 s followed by photobleaching of Cy3 at 15 s.

## Conclusion

In single molecule fluorescence studies, such as smFRET, background fluorescence from labeled substrates often requires their use at concentrations several orders of magnitude less than present *in vivo*. That shortcoming can be addressed through the use of zero-mode waveguides that attenuate background fluorescence by restricting the observation volume. However, the need for expensive and specialized nanofabrication equipment to fabricate ZMWs has precluded their widespread adoption by the biophysical community. Nanosphere lithography allows thousands of ZMWs to be fabricated in parallel, with sizes that are tunable via controlled fusing of polystyrene beads at their glass transition temperature. These robust and inexpensive structures can be made with minimal equipment.

## Materials and Methods

### Slide cleaning

Borosilicate coverslips (No 1.5, Fisherbrand) were cleaned by 10 min sonication in acetone at 40 °C and rinsed three times with distilled water (diH_2_O), followed by a repeat of those steps. They were then sonicated (Branson 5510 ultrasonic bath) in 200 mM KOH for 20 min, rinsed six times with diH_2_O, sonicated in ethanol for 10 min, rinsed three times with diH_2_O, and dried with N_2_.

### Bead deposition

The following protocol describes the formation of 2D crystal lattices of beads on the coverslip surfaces, thus forming the nanomask for colloidal lithography. The cleaned coverslips were each placed in an open, empty petri dish, and the petri dishes were covered by a humidity chamber consisting of an overturned, transparent plastic storage box (76.2 cm x 48.6 cm x 34 cm) along with a small electric fan and a humidity meter (see supporting information, S1 Fig). To prepare beads (Polybead Microspheres 1.00 μm, Polysciences), 200 μL of bead suspension (in water, as provided by the manufacturer) was centrifuged in a 1.5 mL Eppendorf tube for 5 min at 15000 RCF at room temperature, and the water supernatant was removed by pipet. A beaker containing 200 mL of 75 °C H_2_O was then placed inside of the humidity chamber and the fan was turned on, causing the humidity to steadily rise. In the meantime, the beads were resuspended in 100 μL of 1:400 (v/v) TritonX100:ethanol. When the humidity inside of the plastic box reached 85-90%, the lids of the Petri dishes were quickly closed. For each deposition, the plastic box was briefly lifted to allow pipetting of 5 μL of bead suspension into the center of the coverslip, and the lid of the Petri dish was then replaced and the plastic container lowered back down. Each puddle of bead suspension initially spread over a 20 second period, and then receded over the course of 2-3 minutes as the ethanol evaporated, leaving a 2D crystal lattice on the coverslip surface.

### Bead annealing

A flat, milled aluminum plate (13.9 x 13.9 x 0.9 cm), used to provide a uniform temperature surface was placed on top of a hot plate heated to 107 °C (the glass transition temperature of polystyrene), as verified by a thermocouple junction placed on the aluminum plate. When the temperature was stabilized, coverslips with 2D crystal lattices of beads were placed (one-at-a-time) on the surface for 5-25 s and then moved to another aluminum plate at room temperature for rapid cooling.

### ZMW fabrication

The polystyrene mask fabricated as described above underwent thermal or e-beam evaporative deposition (Kurt J. Lesker, PVD75) of either 300 nm Cu (for Al ZMWs) or of 3 nm Ti to serve as an adhesion layer, followed by evaporative deposition of 300 nm of Al (for Au ZMWs). The polystyrene beads were dissolved by soaking in toluene for 2 hours, followed by use of Scotch tape to lift off the metal that had been on top of the beads (rather than on the glass surface). The samples were placed in toluene for another 2 hours, followed by one chloroform and two ethanol rinses. The samples (with either Cu or Al posts) were then oxygen plasma treated for 30 min (Harrick PDC-32G tabletop plasma cleaner), followed by thermal or e-beam deposition of a 3 nm Ti adhesion layer and then 140 nm of either Al or Au cladding. For the Al ZMWs (with Cu posts), the posts were dissolved in copper etchant (copper etchant 49-1, Transene) for one hour, followed by gentle buffing of the surface with lens paper (Thor Labs) to break any remaining posts, and then one more hour in Cu etchant. Similarly, the Al posts were dissolved following one hour in Al etchant (aluminum etchant type A, Transene), a lens paper rub, and one more hour in the Al etchant.

### Scanning probe microscopy

Surface profiling of the metal posts and ZMWs was conducted using a Bruker Bioscope Catalyst AFM in tapping mode, with MikroMasch Opus 160AC-NA standard tips or Nanosensors AR5-NCL high aspect ratio tips.

### Transmission measurements

To obtain the normalized transmission data used in Fig 4, a series of band-pass filters with 10 nm bandwidth were used with the microscope’s illumination tower. The Melles-Griot filters used were 500 nm (03FIV006), 550 nm (03FIV008), 600nm (03FIV018), 650 nm (03FIV002), and 700 nm (03FIV024). The normalized transmission was calculated from intensity data as:

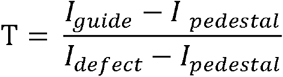

where *I*_*guide*_, is the average intensity of a 2 pixel x 2 pixel square selected to contain the guide emission. *I*_*pedestal*_ is the average intensity of the dark area between guides (see Fig 4). This accounts for the non-zero A/D converter output from the camera even in the dark. *I*_*defect*_ is the average intensity measured from a rectangle placed in the closest defect, a region without any waveguide attenuation.

### Aluminum ZMW functionalization

The following protocol describes the use of orthogonal PVPA chemistry to passivate the Al cladding, and silane chemistry to functionalize the glass bottoms of the wells, as adapted from published protocols [29]. The fabricated ZMWs were first cleaned with a 5 min sonication in acetone, followed by 3 washes with DI water 5 min isopropanol sonication, followed by 3 more washes with DI water. The Al ZMWs were dried with nitrogen and oxygen plasma cleaned for 5 min. A 4.5% (mass/mass) aqueous solution of PVPA was heated to 90 °C. The Al ZMWs were incubated in this preheated solution for 10 min. The ZMWs were then washed with DI water and dried with nitrogen. The ZMWs were finally annealed on a hot plate at 80 °C for 10 min. The Al ZMWs were then treated with 100:5:1 methanol: glacial acetic acid: 3-aminopropyl-triethoxysilane overnight, followed by three diH_2_O rinses, 1 min sonication in EtOH, and were allowed the air dry. The ZMWs then were treated for 3 hrs with 250 mg/mL polyethylene glycol (PEG) in 100 mM NaHCO_3_. 95% of the PEG used was mPEG-succinimidyl valerate (MW 2,000, Laysan), while 5% was biotin-PEG-succinimidyl carbonate (MW 2,000, Laysan). The unbound PEG was rinsed with diH_2_O, followed by drying with N_2_. Functionalized ZMWs were vacuum sealed and stored in the dark at −20 °C until use.

### Gold ZMW functionalization

The following protocol describes the use of orthogonal thiol chemistry to passivate the gold cladding, and silane chemistry to functionalize the glass bottoms of the wells, as adapted from published protocols [30]. The fabricated ZMWs underwent a repeat of the cleaning process used for the initial coverslips (see “Slide Cleaning”), but with no diH_2_O rinse following sonication in ethanol. Instead, they were treated with 100:5:1 methanol: glacial acetic acid: 3-aminopropyl-triethoxysilane overnight, followed by three diH_2_O rinses, 1 min sonication in EtOH, and were allowed to air dry. The samples then were treated for 3 hrs with 250 mg/mL polyethylene glycol (PEG) in 100 mM NaHCO_3_. 95% of the PEG was mPEG-succinimidyl valerate (MW 2,000, Laysan), while 5% was biotin-PEG-succinimidyl carbonate (MW 2,000, Laysan). The unbound PEG was rinsed with diH_2_O, and the slides were dried with N_2_. The samples were then treated overnight with 5 μM mPEG-thiol (MW 2,000, Laysan) in ethanol, before being rinsed and dried again with N_2_. Functionalized ZMWs were vacuum sealed and stored at −20 °C until use.

### Microscopy buffers

Single molecule fluorescence recordings were carried out at 23 °C. For the dsDNA oligos, nuclease-free duplex buffer (IDT) was used, while for ribosomes, TAM_15_ buffer (50 mM Tris-HCl [pH 7.5], 15 mM Mg(OAc)_2_, 30 mM NH_4_Cl, 70 mM KCl, 1 mM DTT) [46] was used. An enzymatic deoxygenation system of 0.3% (w/v) glucose, 300 μg/ml glucose oxidase (Sigma-Aldrich), 120 μg/ml catalase (Roche), and aged 1.5 mM 6-hydroxy-2,5,7,8-tetramethyl-chromane-2-carboxylic acid (Trolox, Sigma-Aldrich) was added to the final single-molecule imaging solutions to reduce fluorophore photobleaching and blinking [46].

### DNA duplex formation

Two complementary single-stranded DNA oligos from Integrated DNA Technologies (IDT) were mixed together at a 1 μM concentration in IDT Duplex Buffer and annealed by placing in a heating block at 95 °C for 1 minute, after which the heating block was turned off and the samples gradually cooled to room temperature and were stored in the dark at 4 °C. Their sequences were as follows:

Biotin,Cy3,Cy5-dsDNA:

CGT GTC GTC GTG CGG CTC CCC AG/iCy3/G CGG CAG TCC
/5Cy5/GGA CTG CCG CCT GGG GAG CCG CAC GAC GAC ACG ACA AAG/3Bio/
Biotin,Cy3-dsDNA:

CGT GTC GTC GTG CGG CTC CCC AG/iCy3/G CGG CAG TCC
GGA CTG CCG CCT GGG GAG CCG CAC GAC GAC ACG ACA AAG/3Bio/
Cy5-dsDNA:

CGT GTC GTC GTG CGG CTC CCC AGG CGG CAG TCC
/5Cy5/GGA CTG CCG CCT GGG GAG CCG CAC GAC GAC ACG ACA AAG
Cy3-dsDNA:

CGT GTC GTC GTG CGG CTC CCC AG/iCy3/G CGG CAG TCC
GGA CTG CCG CCT GGG GAG CCG CAC GAC GAC ACG ACA AAG

### mRNA sequence

5’-biotin-mRNA (Dharmacon, Inc.) had the following sequence, in which the underlined codons form a strong Shine-Delgarno sequence and the italicized codons correspond to the amino acids MFVRYYYYYYYYY:

biotin-AUU UAA AAG UUA <u>AUA AGG AUA</u> CAU ACU *AUG UUC GUG CGU UAU UAU UAU UAU UAU UAU UAU UAU UAU*

### Initiation complex formation

70S initiation complex was formed by incubating 1 μM 70S *E. coli* ribosomes, 4 μM biotin-mRNA (sequence listed above), 1.5 μM each of IF1, IF2, IF3 and fMet-tRNA^fMet^, and 1 mM GTP in TAM_15_ buffer for 25 min at 37 °C.

### tRNAs and ternary complex formation

tRNAs were prepared using the reduction and charging protocols previously described[46,48] starting with *E. coli* tRNA^Val^, and yeast tRNA^Phe^ purchased from Chemical Block (Moscow). tRNA^Phe^ and tRNA^Val^ were conjugated to Cy5 and Cy3 respectively via a dihydroU-hydrazine linkage as described[49]. Ternary complex was formed by incubating 4 μM EF-Tu, 2 μM dye-labeled and charged tRNA, 3 mM GTP, 1.3 mM phosphoenolpyruvate, and 5 μg/mL pyruvate kinase in TAM_15_ buffer for 15 min at 37 °C.

### Flow chamber construction

The sample flow chambers (8 μL) were formed on slides with holes drilled using a 1.25 mm diamond-tipped drill bit. Double-sided tape (Scotch Double Sided, 3M Corporation) laid between the drilled holes served as spacers and separated the flow chambers. Polyethylene tubing with 0.97 mm outer diameter (Warner Instruments) was inserted into each hole and sealed with epoxy (5 Minute Epoxy in DevTube, Devcon). The coverslips with ZMWs were then sealed in place via the double-sided tape and epoxy at their edges. These flow chambers allow fast injection of reaction mixtures into the channels while recording image stacks.

### Fluorescence microscope

A custom-built objective-type total internal reflection fluorescence (TIRF) microscope was used with the incidence angle adjusted to be near TIRF conditions unless otherwise stated. The microscope was based on a commercial inverted microscope (Eclipse Ti, Nikon) with a 1.49 N.A.100× oil immersion objective (Apo TIRF; Nikon). Excitation of Cy3 (and sensitized Cy5 emission during FRET) was achieved with a 532 nm laser, (CrystaLaser, Inc.), while a 640 nm laser (Coherent, Inc.) was used to directly excite Cy5. Fluorescence intensities in a Cy3 channel (550-620 nm) and a Cy5 channel (660-720 nm) were separated by a Dual View splitter (Photometrics) and recorded with an electron-multiplying charge-coupled device (EMCCD) camera (Cascade II, Photometrics) All imaging was conducted at 100 ms frame interval unless otherwise noted.

### DNA duplex imaging in ZMWs

0.5 mg/ml streptavidin was injected for 10 min and washed away with duplex buffer before 100 pM Cy3-Cy5-biotin or Cy3-biotin dsDNA duplex was added and immobilized in the ZMW wells. After 10 min of further incubation, the unbound biotinylated DNA was washed away with buffer and replaced by illumination buffer containing 0 nM to 1 μM of non-biotinylated (background) Cy3-dsDNA or Cy5-dsDNA.

### Ribosome smFRET in ZMWs

0.5 mg/ml streptavidin was injected for 10 min and washed away with TAM_15_ buffer before 100 pM initiation complex was added and immobilized in the ZMW wells via biotin-mRNA. Unbound initiation complex was washed away with TAM_15_ buffer. Solutions containing 10 nM Phe-tRNA^Phe^(Cy5)·EF-Tu·GTP ternary complex, 2 μM EF-G, and 3 mM GTP in TAM_15_ was injected to allow the ribosome to translate through the fMet and Phe codons, forming a POST-translocation state with Phe-tRNA^Phe^ (Cy5) in the P-site. The unbound ternary complex and EF-G were then washed out with TAM_15_ buffer, and the vacant A-sites of the ribosomes were filled by injecting 10 nM Val-tRNA^Val^ (Cy3), forming a stalled PRE-translocation state. The smFRET was then recorded in TAM_15_ imaging buffer with 500 nM added (background) Phe-tRNA^Phe^(Cy5)·EF-Tu·GTP.

### Data processing

Fluorescence traces were extracted from recorded image stacks using using lab-written ImageJ software[46], and analyzed using Matlab. FRET efficiency was calculated according to [50]:

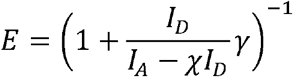

where *I*_*D*_ and *I*_*A*_ are the raw fluorescence intensities of the donor and acceptor above background, *χ* is the cross-talk of the donor emission into the acceptor recording channel, and *γ* accounts for the ratios of quantum yield and detection efficiency between the donor and the acceptor. ImageJ was also used to identify the location of defects in the ZMW array using images of the microscope fields taken with the white light lamp on the illumination tower. The locations of those defects were used to exclude fluorescence traces originating from the defects.

## Acknowledgements

This work was supported by NIH grants R01GM080376 and R35GM118139 to Y.E.G., and by an NIAID pre-doctoral NRSA fellowship F30AI114187 to R.M.J. The manuscript was written through contributions of all authors. All authors have given approval to the final version of the manuscript.

## Supporting Information

**S1 Fig. Homebuilt humidity chamber**

**(a)** Petri dish with lid partially ajar to exchange contents with humid air of the humidity chamber, in which cleaned coverslips are placed before being enclosed in the humidity chamber. **(b)** Layout of several petri dishes (containing coverslips), fan, beaker of water, and humidity monitor on the benchtop prior to covering with the “lid” of humidity chamber. **(c)** Humidity chamber enclosed by the “lid” consisting of an overturned plastic storage box.

**S2 Fig. Schematic of Au ZMW fabrication using nanosphere lithography**.

Black boxes represent expanded insets. T_*g*_ is the glass transition temperature. For Au ZMWs aluminum is used as the post material.

**S3 Fig. AFM characterization of Au devices**.

**(a)** Structural coloration from the diffraction of incident light following Al evaporative coating. **(b)** A representative AFM image of posts formed by Al deposition with the bead template annealed for 20 s. **(c)** Devices after Al deposition and bead lift-off. **(d)** Distribution of Feret diameters of posts formed using a polystyrene bead template annealed for 20 s. **(e)** Completed gold ZMWs. **(f)** A representative AFM image of ZMW wells formed after 25 dissolving the narrow Al posts. **(g)** AFM profile of an individual well. **(h)** Distribution of Feret diameters of wells formed with a bead template that was annealed for 20 s.

**S4 Fig. Example smFRET Traces**

**(A)** Schematic cartoon of biotinylated Cy3-Cy5 dsDNA duplex immobilized in an Al ZMW while illuminated by 532 nm laser light. **(B)** Example smFRET trace of Cy3-Cy5 dsDNA duplex in an Al ZMW. **(C)** Schematic of biotinylated Cy3-Cy5 dsDNA duplex immobilized in an Au ZMW while illuminated by 532 nm laser light. **(D)** Example smFRET trace of Cy3-Cy5 dsDNA duplex in an Au ZMW.

**S5 Fig. Single-Molecule performance with dsDNA at 640 nm excitation**.

**(A)** Schematic cartoon of a biotinylated Cy3-Cy5 dsDNA duplex immobilized in an Al ZMW while illuminated by 640 nm laser light for direct excitation of Cy5, in the presence of non-biotinylated Cy5 dsDNA duplexes in the solution. **(B)** Line histograms depict the distributions of the SNR and **(C)** SBNR of Cy5 emission from immobilized DNA oligos under direct excitation at different concentrations of background Cy5 dsDNA, with bar charts indicating the mean ± SD. **(D)** Example traces of Cy5 emission from direct excitation of immobilized oligos at 0 nM (n =833), 1 nM (638), 10 nM (585), 50 nM (575), 100 nM (238), 500 nM (526), 1000 nM (286) concentrations.

## References

1. Moerner WE, Fromm DP. Methods of single-molecule fluorescence spectroscopy and microscopy. Rev Sci Instrum. 2003;74(8):3597–619.

2. Yildiz A, Ha T, Goldman YE, Selvin PR. Myosin V Walks Hand-Over-Hand□: Single Fluorophore Imaging with 1. 5-nm Localization. Science (80-). 2014;300(5628):2061–6.

3. McKinney SA, Déclais A-C, Lilley DMJ, Ha T. Structural dynamics of individual Holliday junctions. Nat Struct Biol. 2003;10(2):93–7.

4. Forkey JN, Quinlan ME, Alexander Shaw M, Corrie JET, Goldman YE. Three-dimensional structural dynamics of myosin V by single-molecule fluorescence polarization. Nature. 2003;422(6930):399–404.

5. Levene MJ, Korlach J, Turner SW, Foquet M, Craighead HG, Webb WW. Zero-Mode Waveguides for Single-Molecule Analysis at High Concentrations. Science (80-). 2003 Jan;299(5607):682–6.

6. Uemura S, Aitken CE, Korlach J, Flusberg BA, Turner SW, Puglisi JD. Real-time tRNA transit on single translating ribosomes at codon resolution. Nature. 2010 Apr;464(7291):1012–7.

7. Moran-Mirabal JM, Craighead HG. Zero-mode waveguides: sub-wavelength nanostructures for single molecule studies at high concentrations. Methods. 2008 Sep;46(1):11–7.

8. Bethe HA. Theory of Diffraction by Small Holes. Phys Rev. 1944 Oct;66(7–8):163–82.

9. Genet C, Ebbesen TW. Light in tiny holes. Nature. 2007 Jan;445(7123):39–46.

10. Rigneault H, Capoulade J, Dintinger J, Wenger J, Bonod N, Popov E, et al. Enhancement of Single-Molecule Fluorescence Detection in Subwavelength Apertures. Phys Rev Lett. 2005;95(11):117401.

11. Fore S, Yuen Y, Hesselink L, Huser T. Pulsed-Interleaved Excitation FRET Measurements on Single Duplex DNA Molecules Inside C-Shaped Nanoapertures. Nano Lett. 2007;7(6):1749–56.

12. Samiee KT, Foquet M, Guo L, Cox EC, Craighead HG. λ-Repressor Oligomerization Kinetics at High Concentrations Using Fluorescence Correlation Spectroscopy in Zero-Mode Waveguides. Biophys J. 2005;88(3):2145–53.

13. Samiee KT, Moran-Mirabal JM, Cheung YK, Craighead HG. Zero mode waveguides for single-molecule spectroscopy on lipid membranes. Biophys J. 2006;90(9):3288–99.

14. Moran-Mirabal JM, Torres AJ, Samiee KT, Baird BA, Craighead HG. Cell investigation of nanostructures: zero-mode waveguides for plasma membrane studies with single molecule resolution. Nanotechnology. 2007;18(19):195101.

15. Leutenegger M, Gösch M, Perentes A, Hoffmann P, Martin OJF, Lasser T. Confining the sampling volume for Fluorescence Correlation Spectroscopy using a sub-wavelength sized aperture. Opt Express. 2006;14(2):956–69.

16. Perentes A, Utke I, Dwir B, Leutenegger M, Lasser T, Hoffmann P, et al. Fabrication of arrays of sub-wavelength nano-apertures in an optically thick gold layer on glass slides for optical studies. Nanotechnology. 2005 May;16(5):S273.

17. Foquet M, Samiee KT, Kong X, Chauduri BP, Lundquist PM, Turner SW, et al. Improved fabrication of zero-mode waveguides for single-molecule detection. J Appl Phys. 2008;103(3):34301.

18. Fischer UC, Zingsheim HP. Submicroscopic pattern replication with visible light. J Vac Sci Technol. 1981 Nov;19(4):881–5.

19. Deckman HW, Dunsmuir JH. Natural lithography. Appl Phys Lett. 1982 Aug;41(4):377–9.

20. Gang Z, Dayang W. Colloidal Lithography—The Art of Nanochemical Patterning. Chem An Asian J. 2009;4(2):236–45.

21. Wang B, Zhao W, Chen A, Chua S-J. Formation of nanoimprinting mould through use of nanosphere lithography. Vol. 288, Journal of Crystal Growth - J CRYST GROWTH. 2006. 200–204 p.

22. Huang Z, Fang H, Zhu J. Fabrication of Silicon Nanowire Arrays with Controlled Diameter, Length, and Density. Adv Mater. 2007 Feb;19(5):744–8.

23. Tan BJY, Sow CH, Koh TS, Chin KC, Wee ATS, Ong CK. Fabrication of Size-Tunable Gold Nanoparticles Array with Nanosphere Lithography, Reactive Ion Etching, and Thermal Annealing. J Phys Chem B. 2005 Jun;109(22):11100–9.

24. Singh G, Gohri V, Pillai S, Arpanaei A, Foss M, Kingshott P. Large-Area Protein Patterns Generated by Ordered Binary Colloidal Assemblies as Templates. ACS Nano. 2011;5(5):3542–51.

25. Singh G, Pillai S, Arpanaei A, Kingshott P. Highly Ordered Mixed Protein Patterns Over Large Areas from Self-Assembly of Binary Colloids. Adv Mater. 2011;23(13):1519–23.

26. Willets KA, Van Duyne RP. Localized Surface Plasmon Resonance Spectroscopy and Sensing. Annu Rev Phys Chem. 2007;58(1):267–97.

27. Hulteen JC, Van Duyne RP. Nanosphere lithography: A materials general fabrication process for periodic particle array surfaces. J Vac Sci Technol A. 1995 May;13(3):1553–8.

28. Haynes CL, McFarland AD, Smith MT, Hulteen JC, Van Duyne RP. Angle-Resolved Nanosphere Lithography: Manipulation of Nanoparticle Size, Shape, and Interparticle Spacing. J Phys Chem B. 2002 Feb;106(8):1898–902.

29. Iwasa T, Han Y-W, Hiramatsu R, Yokota H, Nakao K, Yokokawa R, et al. Synergistic effect of ATP for RuvA–RuvB–Holliday junction DNA complex formation. Sci Rep. 2015 Dec;5:18177.

30. Kinz-Thompson CD, Palma M, Pulukkunat DK, Chenet D, Hone J, Wind SJ, et al. Robustly Passivated, Gold Nanoaperture Arrays for Single-Molecule Fluorescence Microscopy. ACS Nano. 2013 Sep;7(9):8158–66.

31. Denkov N, Velev O, Kralchevski P, Ivanov I, Yoshimura H, Nagayama K. Mechanism of formation of two-dimensional crystals from latex particles on substrates. Langmuir. 1992 Dec;8(12):3183–90.

32. Cheung CL, Nikolic RJ, Reinhardt CE, Wang TF. Fabrication of nanopillars by nanosphere lithography. Nanotechnology. 2006;17(5):1339–43.

33. Peng K, Zhang M, Lu A, Wong N-B, Zhang R, Lee S-T. Ordered silicon nanowire arrays via nanosphere lithography and metal-induced etching. Appl Phys Lett. 2007;90(16):163123.

34. Micheletto R, Fukuda H, Ohtsu M. A Simple Method for the Production of a Two-Dimensional, Ordered Array of Small Latex Particles. Langmuir. 1995 Sep;11(9):3333–6.

35. Vukusic P, Sambles JR. Photonic structures in biology. Nature. 2003 Aug;424(6950):852–5.

36. Ball P. Nature’s Color Tricks. Sci Am. 2012;306(5):74–9.

37. Rieger J. The glass transition temperature of polystyrene. J Therm Anal. 1996;46(3):965–72.

38. Kosiorek A, Kandulski A, Glaczynska H, Giersig M. Fabrication of Nanoscale Rings, Dots, and Rods by Combining Shadow Nanosphere Lithography and Annealed Polystyrene Nanosphere Masks. Small. 2005 Feb;1(4):439–44.

39. Hanarp P, Käll M, Sutherland DS. Optical Properties of Short Range Ordered Arrays of Nanometer Gold Disks Prepared by Colloidal Lithography. J Phys Chem B. 2003 Jun;107(24):5768–72.

40. Fujimoto K, Morita Y, Iino R, Tomishige M, Shintaku H, Kotera H, et al. Simultaneous Observation of Kinesin-Driven Microtubule Motility and Binding of Adenosine Triphosphate Using Linear Zero-Mode Waveguides. ACS Nano. 2018;12(12):11975–85.

41. Degiron A, Lezec HJ, Yamamoto N, Ebbesen TW. Optical transmission properties of a single subwavelength aperture in a real metal. Opt Commun. 2004 Sep;239(1–3):61–6.

42. Shroder DY, Lippert LG, Goldman YE. Single molecule optical measurements of orientation and rotations of biological macromolecules. Methods Appl Fluoresc. 2016;4(4):042004.

43. Roy R, Hohng S, Ha T. A practical guide to single-molecule FRET. Nat Methods. 2008;5(6):507–16.

44. Pollack G, Daniel S. Electromagnetism. 1st ed. Addison-Wesley; 2001.

45. McCann JJ, Choi UB, Zheng L, Weninger K, Bowen ME. Optimizing Methods to Recover Absolute FRET Efficiency from Immobilized Single Molecules. Biophys J. 2010 Aug;99(3):961–70.

46. Chen C, Stevens B, Kaur J, Cabral D, Liu H, Wang Y, et al. Single-molecule fluorescence measurements of ribosomal translocation dynamics. Mol Cell. 2011 May;42(3):367–77.

47. Jamiolkowski RM, Chen C, Cooperman BS, Goldman YE. tRNA Fluctuations Observed on Stalled Ribosomes Are Suppressed during Ongoing Protein Synthesis. Biophys J. 2017 Dec;113(11):2326–35.

48. Pan D, Qin H, Cooperman BS. Synthesis and functional activity of tRNAs labeled with fluorescent hydrazides in the D-loop. RNA. 2009 Feb;15(2):346–54.

49. Kaur J, Raj M, Cooperman BS. Fluorescent labeling of tRNA dihydrouridine residues□: Mechanism and distribution. RNA. 2011;7:1393–400.

50. Ha T, Ting AY, Liang J, Caldwell WB, Deniz AA, Chemla DS, et al. Single-molecule fluorescence spectroscopy of enzyme conformational dynamics and cleavage mechanism. Proc Natl Acad Sci U S A. 1999 Feb;96(3):893–8.

