## Supplementary figures and images for "Nanoaperture fabrication via colloidal lithography for single molecule fluorescence imaging"

### Supplemental Fig 1

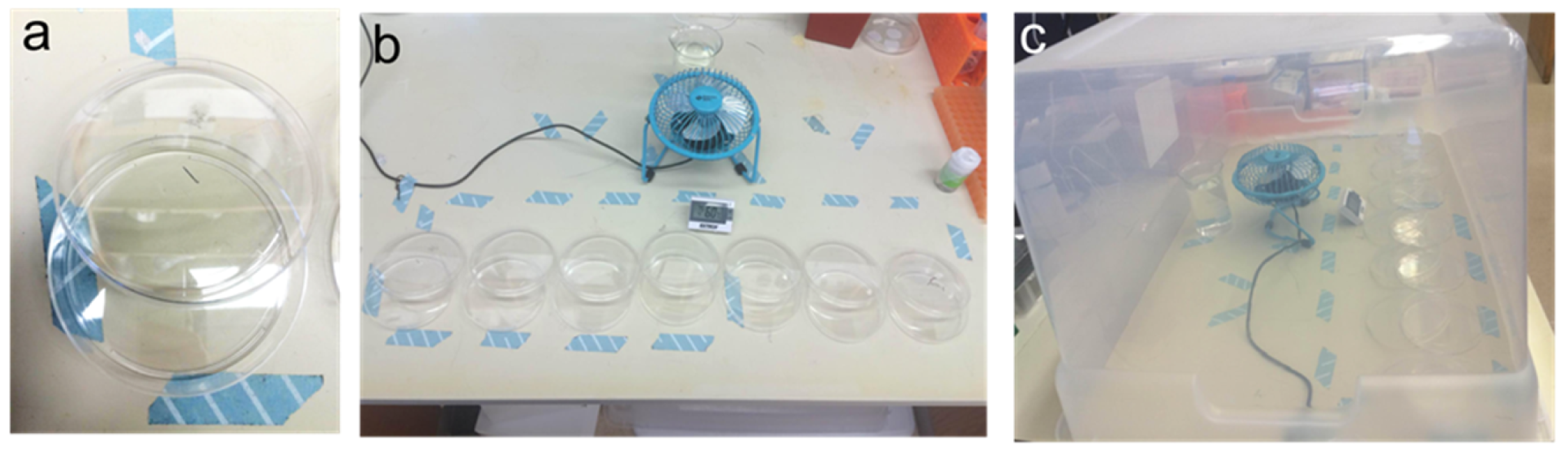

### Supplemental Fig 2

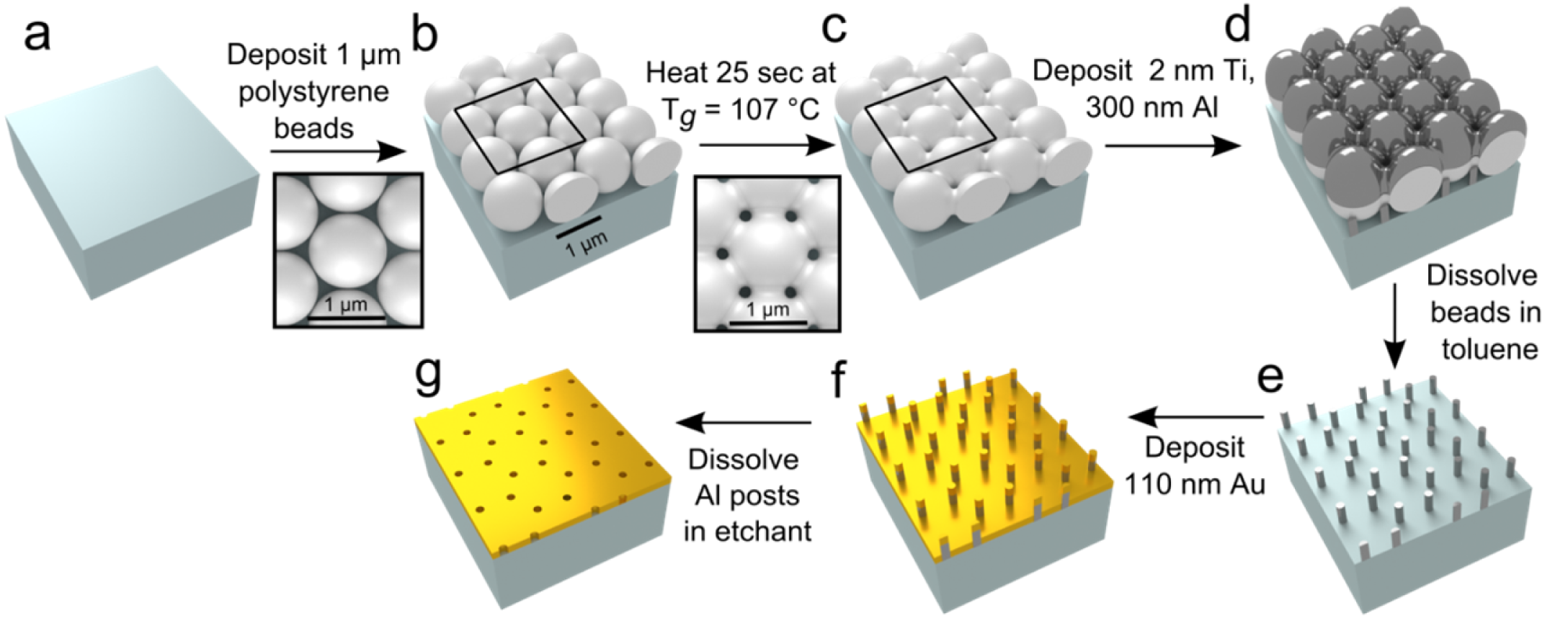

### Supplemental Fig 3

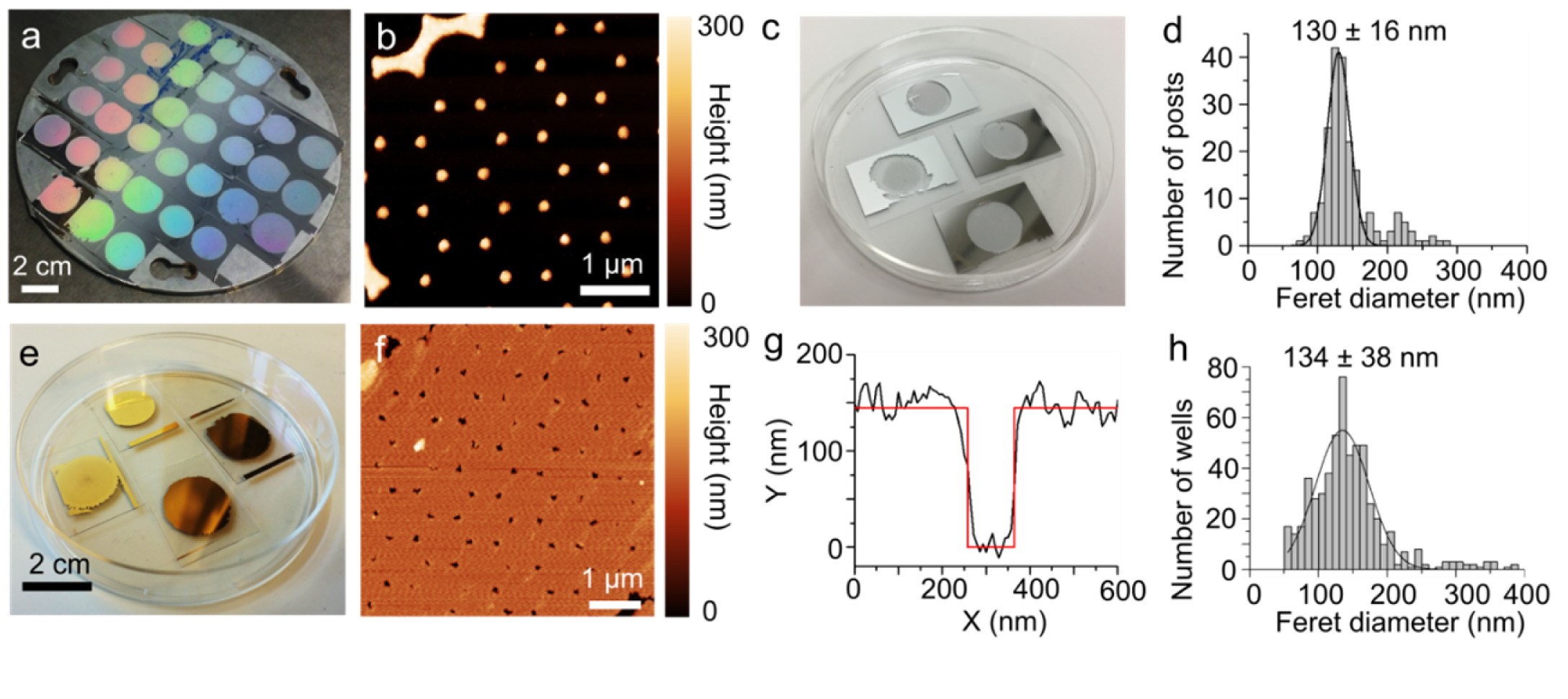

### Supplemental Fig 4

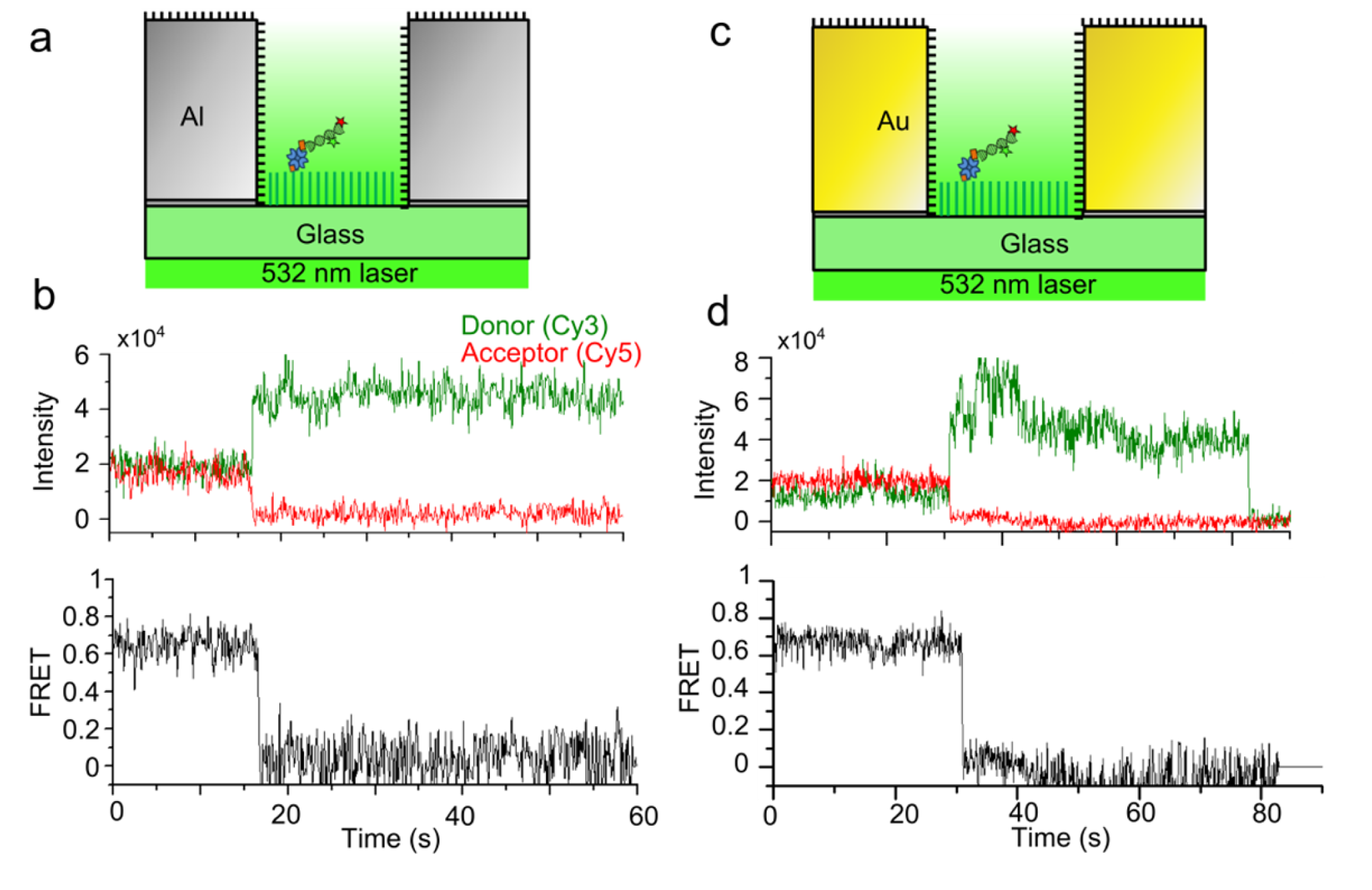

### Supplemental Fig 5

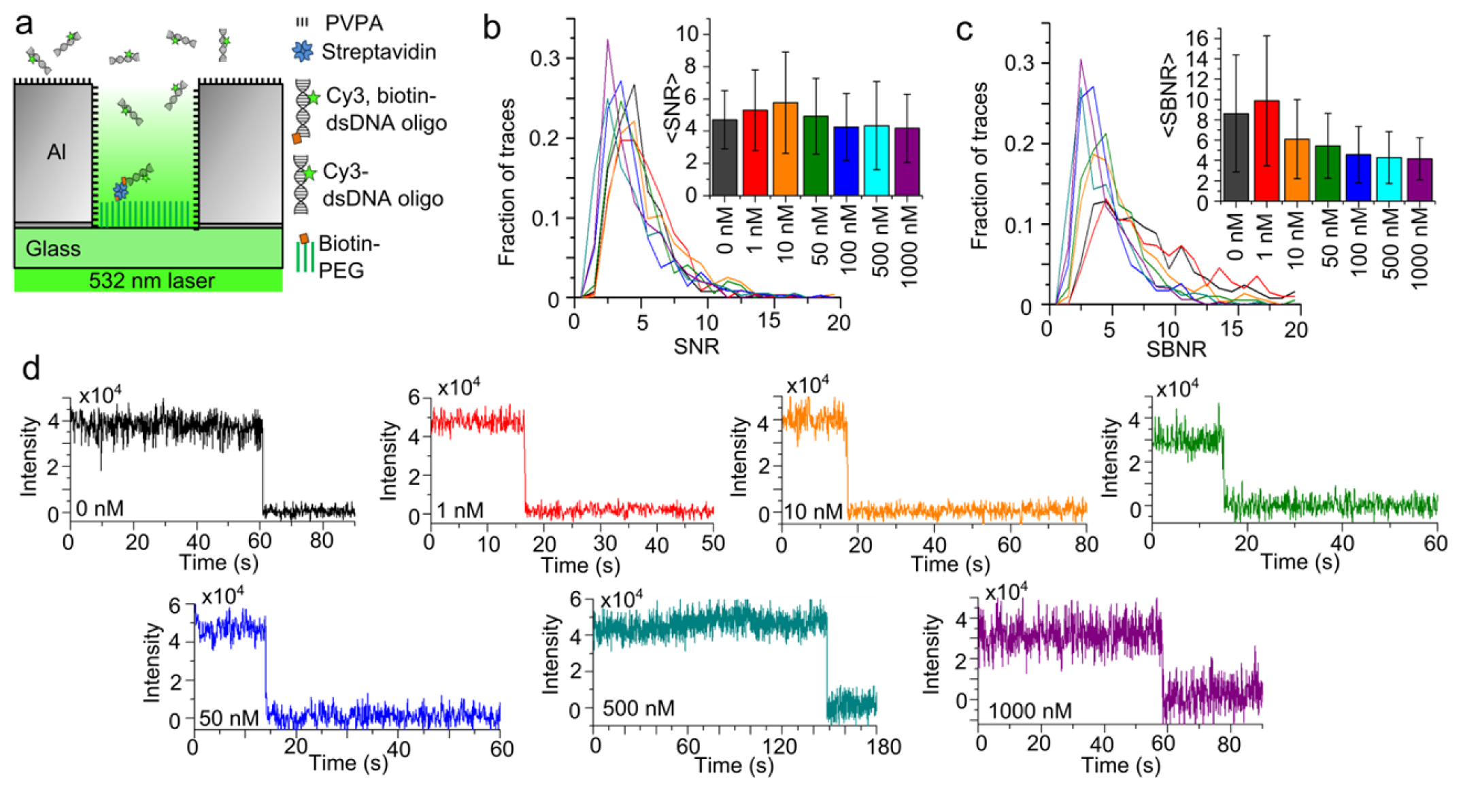
